# Zmap: an intelligent region-allocation method to map single-cell into spatial data

**DOI:** 10.1101/2025.01.27.635178

**Authors:** Quanyou Cai, Lihui Lin, Xin Liu, Jiekai Chen

## Abstract

Integration of single-cell and spatial transcriptome represents as a fundamental strategy to enhance spatial data quality. However, existing methods for mapping single-cell data to spatial coordinates struggle with large-scale datasets comprising millions of cells. Here, we introduce Zmap, an intelligent region-allocation method inspired by the region-of-interest (ROI) concept from image processing. By using gradient descent, Zmap allocates cells to structured spatial regions that matching the most significant biological information, optimizing the integration of data and improving speed and accuracy. Zmap excels in integrating data even in the presence of various sequencing artifacts, such as cell segmentation errors and imbalanced cell-type representations. Zmap outperforms state-of-the-art methods by 10 to 1000 times in speed, and it is the only approach capable of integrating datasets containing millions of cells in a single run. As a result, Zmap uncovers originally hidden gene expression patterns in the brain section, offers new insights into organogenesis and tumor microenvironments, all with exceptional efficiency and accuracy.

## INTRODUCTION

Cells in a multicellular organism must be able to signal and respond to both internal and external spatial cues. Understanding how the cellular blueprint unfolds into tissues and organs is key to revealing how cells differentiate, communicate and organize(1,2). Even high-resolution spatial transcriptome technologies have been developed, cell segmented data from such technologies showed significantly less discrimination compared to single-cell transcriptome data. Therefore, integrating single-cell transcriptome and spatial transcriptome is important for understanding development and disease(3).

Data integrating allows scRNA-seq data to acquire spatial coordinates while increasing the number of detected genes in the spatial space(4–6), and has improved the resolution of analyzing spatial cell heterogeneity and niche in developmental and disease tissues (7–9). However, the mismatch in single cell and spatial data caused by sequencing noise and sampling bias hinders accurate spatial mapping. For example, cell segmentation errors can lead to distortion of spatial cell gene expression and confound cell identity (10). And even for spatial transcriptome and single cell sequencing data from the same tissue source, the proportions of cell types are not the same(11,12).

To address these challenges, we developed Zmap, a framework that utilizes multiple layers of regionalization constraints to localize the cells’ position with high noise tolerance and computational efficiency. We demonstrate that Zmap precisely localizes the single cells, thereby reconstructing the correct spatial gene expression pattern, improving the classification ability of spatial data, and enhancing the inference of inter-cellular interactions. Furthermore, this user-friendly tool accommodates various customized regionalization patterns, making it adaptable to complex tissue and organ structures. By integrating spatial and single-cell omics data, Zmap deepens our understanding of cell lineages and spatial dynamics in tissue and organ development, offering valuable insights into both physiological and pathological processes.

## MATERIAL AND METHODS

### Zmap algorithm overview

Given a single-cell gene expression profile *S* with dimensions *n*_*cells*_ × *n*_*genes*_, and a ST gene expression *T* profile with dimensions *n*_*voxels*_ × *n*_*genes*_, the goal of Zmap is to map cells in scRNA-seq data to spatial voxels in ST data. The voxels could be spatial cells predicted by image segmentation, circular spots or bins comprised of spots. The algorithm can be used for both imaging-based and sequencing-based ST data. Both *S* and *T* should be non-normalized count matrices for the calculation, such that *S*_*ik*_ is the expression count of gene *k* in cell *i*, and *T*_*vk*_ is the expression count of gene *k* in voxel *v*. The data integration process consists of four steps.

#### Gridding

The integration begins by gridding the ST data, which is divided into a number bins of equal-sized square (or hexagons), where the bins are connected edge-to-edge. Each spatial cell is assigned to a bin based on the position of its nucleus coordinate, and the expression values of the cells within each bin are summed to form matrix *G* of dimensions *n*_grids_ × *n*_genes_, where *G*_*qk*_ is the expression level of gene *k* in grid *q*. One of advantages of gridding is to improve the signal-to-noise ratio of ST data. Cell segmentation errors are likely to occur within a tissue with a large number of tightly connected cells. In this scenario, the predicted boundaries of each cell differ from their actual boundaries. This discrepancy results in differences between the spatial gene expression and true gene expression of cells. However, when we group neighboring cells together by gridding, the effects of segmentation errors only influence the cells at the edges of the bin, without impacting the cells inside the bin. This principle remains applicable when aggregating multiple bins into larger spatial regions. The other advantage of gridding is to reduce the cost of mapping calculation. Zmap first maps cells to bins followed by voxels within each bin, instead of maps cells to all voxel directly. Since the number of bins is much less than the number of spatial voxels, this strategy enables the algorithm to integrate large-scale data. We suggest that users adjust the size of the bin to cover 3 - 15 voxel (Specially, 1 voxel for 10x Visium), Zmap provides bar plot to count the number of cells in the grids. Within this range, different bin sizes have little impact on the results of data integration.

#### Spatial Regionalization

Zmap combines the bins into horizontal and vertical stripes, representing two default layers of regionalization. Similarly, for ST data like 10x Visium, each spot is considered as a hexagon, which can be combined into three direction stripes with a 60° angle, representing three layers of regionalization. The combination of these stripes enables the positioning of all bins. Zmap also uses the SpaGCN(13) algorithm to cluster bins, generating another layer of regionalization. Zmap is a flexible framework that allows users to input custom layers of regionalization, such as physiological and pathological regions defined by tissue imaging, or clusters generated by other clustering methods.

For a layer *l* of regionalization, we define an *n*_*regions*_ × *n*_*bins*_ one-hot encoding matrix *P*^(*l*)^ to represent the region label of all bins, where 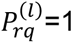 denotes the bin *q* belong to region *r*. We define region gene expression matrix *R*^(*l*)^ with dimension of *n*_*regions*_ × *n*_*genes*_, where 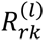 is the sum expression of all spatial voxels in region *r*.

#### Mapping cells to bins

We aim to solve the mapping matrix *M* with dimension *n*_*cells*_ × *n*_*bins*_ by minimizing the dissimilarity between the sum expression of single cells assigned to each region, and the sum expression of spatial voxels in each region. The object function is formulated as follows:

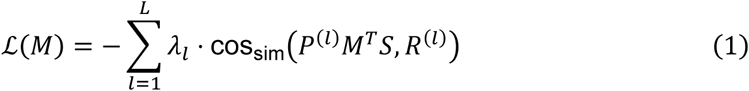

Where cos_sim_ is the cosine similarity function 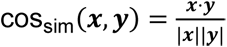, *L* is the total number of layers, and *λ*_*l*_ is the weight of each layer, by default *λ*_*l*_ = 1 for all layers. The mapping matrix *M* is converted to a probability matrix using the softmax function:

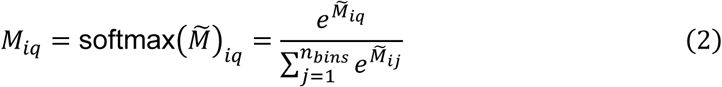

The softmax function ensures that 0 ≤ *M_iq_* ≤ 1 and 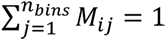.

The optimal *M* is obtained by Adam gradient descent algorithm in the PyTorch library. Zmap then filters out cells with mapping probabilities less than threshold *t*, we set *t* = 0.1 as the default value. Each of the remaining cells can be assigned to one or more bins.

Generally, as the number of regionalization layers *L* increases, the constraints on cell localization become more accurate, and the computation time will also increase. In our experiments, we found that utilizing only 3 to 4 layers constraints is enough to achieve highly accurate cell localization.

#### Mapping cells to voxels

Within each bin, Zmap calculates the cosine similarity between spatial voxels and single cells. The cell with the maximum cosine similarity value is selected for each voxel. In this process, a single cell can be assigned to multiple voxels. We also provide a filtering parameter *c* to remove cell mappings with low similarity.

### Benchmarking analysis with real adjacent sections datasets

To evaluate the performance of Zmap, we conducted benchmarking analysis using single-cell resolution data from adjacent sections of an E8.5 mouse embryo(14), generated through the SeqFISH technique. The data were obtained from two sections of the same embryo, spaced 12 mm apart (labeled as L1 and L2). L1 data (10,150 cells) were used as the spatial dataset, while L2 data (7,656 cells) were used as the single-cell dataset for spatial reconstruction.

Similarly, we evaluated another spatial transcriptomics technique, STARMAP(15), using mouse cortical data obtained from two adjacent sections. One section (8,506 cells) was used as the spatial dataset, while the other (9,803 cells) served as the single-cell dataset for reconstruction. This allowed us to benchmark Zmap’s performance across different spatial transcriptomics platforms and datasets.

### Metrics for benchmarking performance

To quantitatively evaluate the performance of spatial reconstruction, we employed the following metrics, each assessing a specific aspect of accuracy or similarity between true and predicted data.

#### Location Error

We calculated the mean absolute error (MAE) of reconstructed spatial coordinates to measure spatial accuracy. For each cell, the Euclidean distance between its true and predicted coordinates was computed. The MAE for each cell type was then derived as:

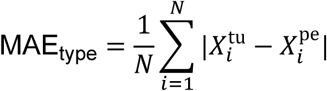

Where 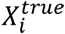 and 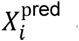 are the true and predicted coordinates of cell i, and N is the total number of cells of a given type.

#### Hit Number (Neighborhood Overlap)

We evaluated local spatial consistency using the overlap of nearest neighbors between the true and reconstructed coordinates. For each cell, we identified its k-nearest neighbors (*k* = 5,10,…,50) in both datasets and computed the intersection of these neighbor sets. The mean hit number for each k is given by:

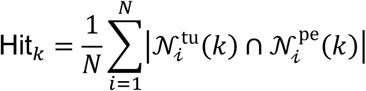

Where 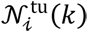 and 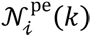 are the sets of true and predicted neighbors for cell i. We utilized PCC-voxel to study the similarity in spatial distribution patterns of cell types between real spatial transcriptomics data and spatially reconstructed single-cell data. To achieve this, we encoded a binary variable (0 or 1) for each real spatial location, indicating whether the location is occupied by a specified cell type. The Pearson correlation coefficient for the cell type distributions between the real and reconstructed data was calculated. Otherwise, to evaluate spatial alignment across the dataset, we divided the spatial domain into grids of fixed size. For each grid, the frequency of cells of each type was calculated. We then compared these frequencies between the true and predicted data using Pearson correlation:

### Simulations of spatial detection noise

For cell segmentation noise, we simulated a ground truth ST data with real single cel. Each spatial cell in the seqFISH+ data(16) was replaced by a single cell from the snRNA-seq dataset (17) derived from the same tissue. Let *G*_ST_ ∩ *G*_sc_ represent the overlapping genes between the two datasets. For each cell type *c* present in the ST data, subsets 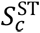 *and* 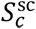 were created, ensuring that 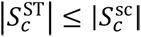. If 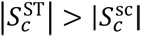, the ST subset was randomly downsampled to size 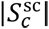. To compute cell similarity, the cosine distance 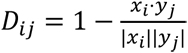 was calculated for each pair of expression profiles 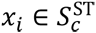 *and* 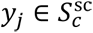, resulting in a distance matrix *D*. The Hungarian algorithm was applied to *D* to find the optimal one-to-one mapping of spatial cells to single cells, producing a set of matches *G* = {(*i*, *j*)}. Spatial coordinates from the ST data were then assigned to the matched single-cell data, generating a single-cell data *A*^sc^ that preserved the spatial structure of the original ST data while incorporating segmentation noise. Next, variability in the simulated ST data was introduced by randomly splitting and redistributing gene expression profiles across neighboring cells. For each cell *k*, its expression *e*_*k*_ was divided into *n* ∈ {2, 3, 4} bins based on normalized percentages *p* = (*p*_1_, *p*_2_,…, *p*_*n*_), ∑ *p*_*i*_ = 1, resulting in bins *b*_*i*_ = *p*_*i*_ ⋅ *e*_*k*_. Neighboring cells *N*(*k*) were identified based on spatial proximity, defined by a maximum spatial distance *d*_max_, default is 250. Expression bins were redistributed to 1–2 randomly selected cells *j* ∈ *N*(*k*), ensuring *e*_*j*_ + *b*_*i*_ ≥ 0. If *N*(*k*) = Ø, redistribution for *k* was skipped. This process introduced controlled noise and variability into the data, enabling robust testing of spatial transcriptomics algorithms under simulated conditions.

### Simulations of cell proportions

We simulated three different scenarios of cell proportion interference: Scenario 1, We used seqFISH+ data(16) as the ground truth for spatial transcriptomics and obtained simulated single-cell data with different cell type proportions by randomly sampling this data. Scenario 2, We used seqFISH+ data as the ground truth for spatial transcriptomics and obtained simulated single-cell data with different cell type proportions, excluding the ExcitatoryL2and3 cell type, by randomly sampling the data. Scenario 3, We used seqFISH+ data with the ExcitatoryL2and3 cell type removed as the ground truth for spatial transcriptomics and obtained simulated single-cell data with different cell type proportions, including one additional cell type compared to the spatial data, by randomly sampling the original data. We evaluated the results using the Euclidean distance between the true spatial locations of the cells and their positions after allocating with the algorithm.

### Mapping single cell data to in situ sequencing spatial data

For STARMAP data(18), we performed spatial mapping of processed SMART-Seq2 snRNA-seq data(17), consisting of 14,249 cells and 34,041 genes, onto 1,523 spatial cells. Since STARMAP detected only 981 genes, we used the shared genes for mapping.

We evaluated the accuracy of the mapping method by calculating spatial distribution similarity of cell types between real spatial transcriptomics data and reconstructed single-cell data. Additionally, we compared the cell type proportions of the reconstructed cells with the proportions measured by STARMAP. The algorithm integrates three layers of information: two layers of regular horizontal and vertical regionalization, and one layer of spatial grid clustering information. The Zmap algorithm converged after 500 epochs with default parameters.

### Mapping single cell data to smFISH-based spatial data

For seqFISH data(14), we performed spatial mapping of processed 10X v2 scRNA-seq data(14), consisting of 116,312 cells and 29,452 genes, onto 19,416 spatial cells. Since seqFISH detected only 351 genes, we used the shared genes for mapping. We evaluated the accuracy of the mapping method by calculating ST-NNED. The algorithm integrates three layers of information: two layers of regular horizontal and vertical regional segmentation, and one layer of spatial grid clustering information. The Zmap algorithm converged after 500 epochs with default parameters.

### Mapping single cell data to Visium data

We used processed data of colorectal cancer-derived liver metastatic tumor (19) from the BD Rhapsody Single-Cell Whole Transcriptome platform, sequencing a total of 115,818 cells across 16,478 genes, which were then mapped to Visium data of same cancer tissue, consisting of 4,672 spots and 17,934 genes. For mapping onto the Visium data, we integrated information from four layers: three directional layers at regular angles (0 degrees, 60 degrees, and 120 degrees) for regional segmentation, and one layer containing the spatial clustering information of the Visium spots. The Zmap algorithm converged after 500 epochs with default parameters. Since the Visium data does not have single-cell resolution, after mapping cells to spots, we assigned each cell to the spot with the highest mapping probability. To visually represent this, we added random jitter to each cell’s coordinates based on the spot’s coordinates, confined within the spot’s radius, to account for multiple cells within the same spot. We used the Milo package to assess the distribution tendency of single cells between the PT and T zones, using an absolute log fold change greater than 2 as the threshold to distinguish cells that prefer different zones. These cells were then subjected to differential analysis.

### Mapping single cell data to Stereo-seq data

We used processed 10X v3 scRNA-seq data of E16.5 mouse embryo, consisting of 169,307 cells and 55,450 genes, mapped to processed stereo-seq data from MOSTA(20). The spatial data used in this study originated from E16.5 mouse embryo samples, which were divided into five sections. We used the results from section 3 for demonstration. This section contained 155,741 spots and 28,579 genes. The algorithm employed a grid with a width of 5 for computation, incorporating three layers of regional constraints: regular horizontal and vertical regional information and grid clustering information. Convergence was achieved after 500 epochs. To accelerate computation, we randomly split the single-cell data into five subsets based on cell types and performed mapping on each subset separately.

### Compared methods

We compared Zmap with five state-of-the-art methods: Tangram (version 1.0.4) (21), scSpace (version 1.0.0)(22), SpaTrio (version 1.0.0)(23), CytoSpace (version 1.0.6a0) (24) and CelEry (version 1.2.1)(25). Tangram optimally maps single cells to spatial voxels primarily by minimizing the cosine distance between the reconstructed single-cell data and spatial voxels. It achieves high performance in aligning single-cell data with spatial transcriptomics, as demonstrated in a previous benchmark study, ensuring a detailed and contextually relevant representation of cellular distributions. We used all genes for calculations and set the parameters to default. scSpace employs transfer learning with Transfer Component Analysis to co-embed scRNA-seq and spatial transcriptomics data into a shared latent space. It reconstructs a pseudo-space for scRNA-seq data by training a multi-layer perceptron on spatial coordinates derived from spatial transcriptomics. Additionally, scSpace introduces spatially informed clustering to identify spatially heterogeneous cell subpopulations. We ran scSpace with its default parameters. SpaTrio integrates single-cell multi-omics and spatial transcriptomics data through fused Gromov-Wasserstein optimal transport. By leveraging gene expression and spatial topology, it reconstructs multimodal tissue maps and identifies spatial co-expression patterns. SpaTrio provides probabilistic cell-spot alignments, enabling accurate mapping of cellular modalities in complex tissues. We used SpaTrio with its default parameters for all analyses. CytoSpace estimates cell types in spatial data through deconvolution before linearly assigning cells to voxels to maximize the Pearson correlation coefficient (PCC). In this study, we ran CytoSpace with the following parameters: cytospace -sc -sp ./sc_exp.tsv -ctp ./sc_anno.tsv - stp ./st_exp.tsv -cp ./st_coor.tsv -nop 4 -noss 1000 -sm lap_CSPR. CelEry applies a supervised deep learning framework to recover spatial origins of cells in scRNA-seq data by learning gene expression spatial location relationships from spatial transcriptomics. It includes an optional data augmentation step using variational autoencoders, enhancing robustness and reducing noise in single-cell data. For CelEry, we set num_workers=10 and used default values for all other parameters

### Computation time and resource analysis

We evaluated the computation time and resource usage using stereo-seq data and corresponding single-cell data. Both the stereo-seq and single-cell data were randomly downsampled to generate spatial data and single-cell data with varying numbers of spots, and we used 25,059 shared genes. We then performed mapping using Zmap (default parameters, width of 5, three layers of region information constraints), Tangram (default parameters, cells mode), CytoSpace (default parameters, sc mode, noss set to 1000, solver using lap_CSPR), scSpace (default parameters), SpaTrio (default parameters), and CelEry (default parameters, with num_workers set to 10). The computation time and resource usage for each method were recorded.

All calculations were tested on the same system, equipped with an Intel E5-2650 v3 (2.3 GHz base and 3.0 GHz max frequencies, 40 cores, and 128GB RAM) and a 24GB NVIDIA GeForce RTX 3090.

## RESULTS

### The algorithm design of Zmap

Zmap begins by creating square (or hexagon) bins that tile the entire spatial profile containing capture area, each bin contains several spatial voxels. By leveraging the tissue architecture, spatial gene expression or virtual generated structure, the bins are organized into diverse layers of regionalization, e.g. virtually generated horizontal stripes, vertical stripes, and spatial clustering domains. Zmap first maps cells to bins. In this process, the different regionalization layers are used to jointly constrain the spatial localization of single cells, and each cell is assigned to the bins with the highest mapping probability. Given the significantly lower number of bins compared to spatial voxels, this approach reduces the computational parameters necessary for calculating the cell-cell mapping probability. Consequently, Zmap is effective for integrating large datasets. Next, within each bin, Zmap pairs the single cell and the spatial cell with the highest similarity, and the spatial voxels are replaced by single cells for downstream analysis (Figure 1A).

**Figure 1:**
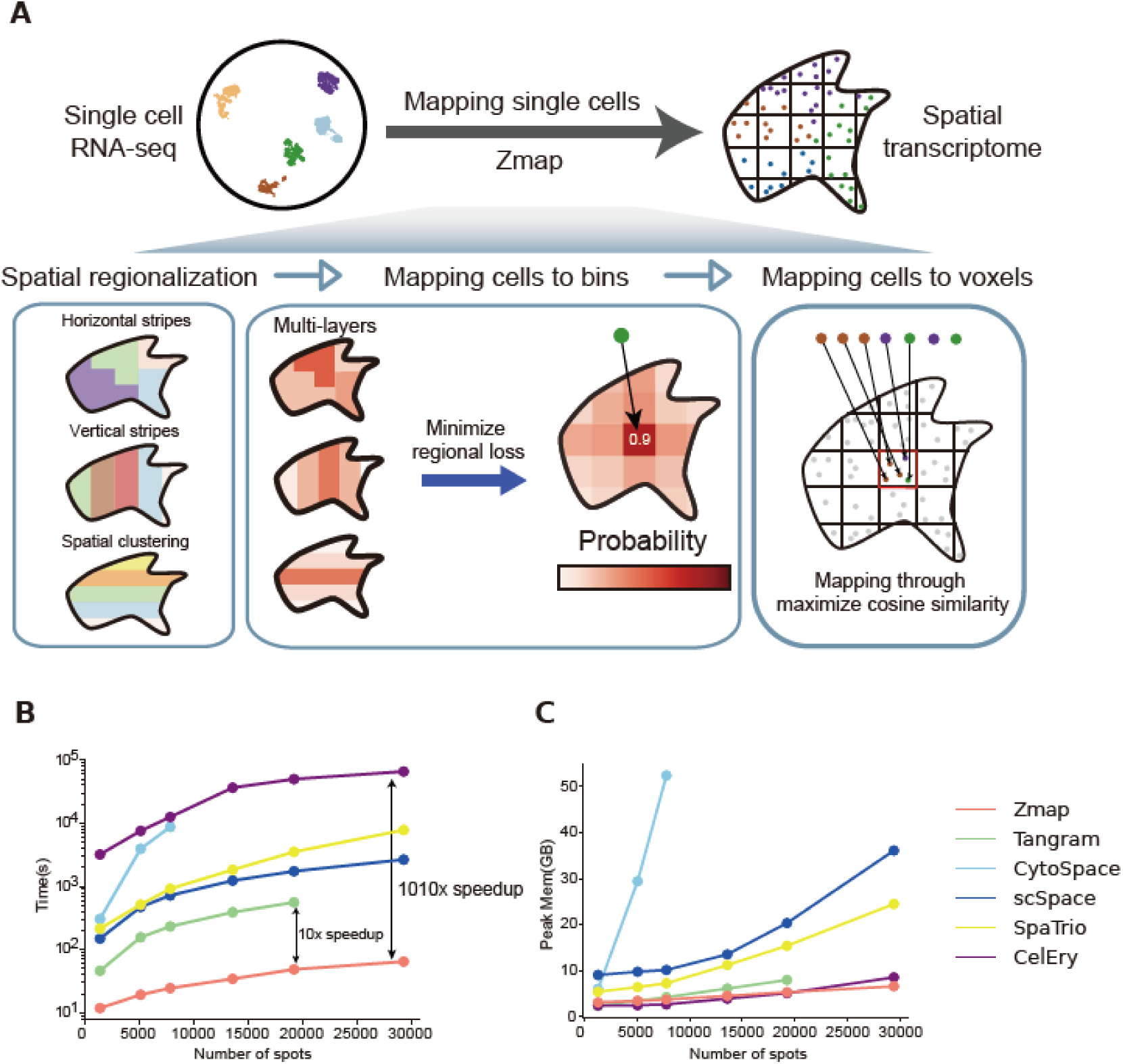
Schematic overview of Zmap. (**A**) Zmap is designed to accurately assign single-cell locations. Starting with spatial region expression formed from gridding the original ST data and single-cell seq data(left), Zmap uses multi-layers regional constraints to optimize the distribution of each cell in each grid(middle), and finally determining the precise location of cells within each grid through min cosine similarity cost(right). (**B**) Computation time of different methods when integrating 5k single cells with varying voxel numbers. (**C**) Peak memory of different methods when integrating 5k single cells with varying voxel numbers.

We used seqFISH+ data of cerebral cortex(16) to illustrate the advantage of multi-layer spatial constraints strategy on mapping spatial location of cells. The dataset contains 10,000 detected genes and 523 spatial cells. We simulated single cell data by adding Gaussian noise to the spatial cells. We then used Zmap to integrate spatial cells and simulated single cells. The known spatial localization at a cellular level in the seqFISH+ data serves as the ground truth for predicting the spatial localization of individual cells. Applying a single layer of constraint resulted in few exact matched cells. Two layers of constraints improved the exact match ratio to ∼50%. When three layers of constraints were applied, the exact match ratio dramatically increases and reaches 100% (Supplementary Figure S1A-S1B). These results highlight significant improvement by importing additional spatial context to localize single cells.

Due to the algorithmic design of Zmap, it demonstrates advantages in runtime and memory usage. We fixed the number of spatial cells, and showed that the cost of time increased by the number of single cells for Zmap and five compared methods (Tangram, scSpace, SpaTrio, CytoSpace and CelEry). The time consumption of Zmap is well controlled for dataset size of both single cells and spatial cells, with the task of integrating 15,000 cells and 19,230 spots taking only 2.4 minutes to complete. Tangram takes more than 24 minutes to complete this task, whereas other methods require longer time, and CytoSpace being unable to complete the task (Figure 1B). Even when the cell and spot number are reduced to 5,000 and 7,829, respectively, CytoSpace still takes more than 144 minutes to finish (Figure 1B and Supplementary Table 1). In addition, the memory usage of Zmap, CelEry and Tangram is comparable and much smaller than others (Figure 1C and Supplementary Table 1). Zmap accurately mapped single cells to the organ regions such as brain, lung, and kidney (Supplementary Figure S2), while maintaining high computational efficiency. These results highlight the exceptional efficiency and scalability of Zmap, making it a robust choice for integrating large-scale spatial and single-cell transcriptomic datasets.

### Benchmarking analysis with adjacent sections datasets

To benchmarking the performance of Zmap in cell spatial reconstruction, we utilized single-cell-resolution mouse embryo data generated by SeqFISH. The dataset includes two sections from distinct tissue layers of the same E8.5 mouse embryo, (designated as L1 and L2). We used the L1 layer (containing 10,150 cells) as the reference spatial dataset and the L2 layer as the single-cell dataset (Figure 2A). We compared five mapping methods using evaluation metrics such as mean absolute error (MAE) of location difference between true and reconstructed coordinates, the spatial distribution similarity of cell types, and the number of true neighbors among the k nearest neighbors for each cell.

**Figure 2:**
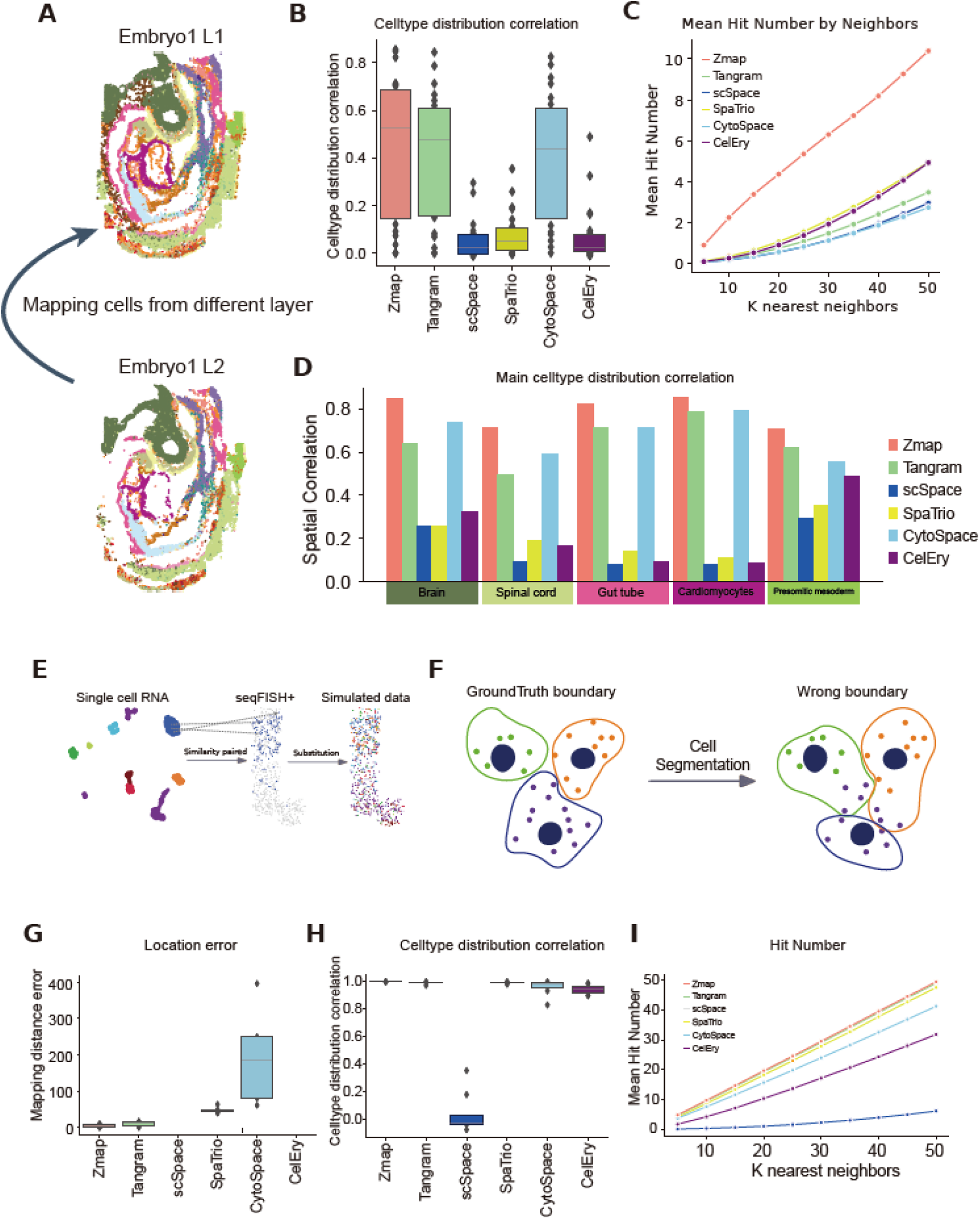
Benchmark analysis. (**A**) Schematic diagram of cell reconstruction using adjacent slices. (**B**) Evaluation of spatial distribution similarity of cell types using different methods. (**C**) Assessment of the number of true neighboring cells. (**D**) Evaluation of spatial distribution similarity for five major cell types. (**E**) Schematic diagram for generating simulated single-cell spatial data. We utilize seqFISH+ data and single-cell data from similar tissues, extract shared cell types, and for each cell in the spatial data, identify all corresponding cells of the same type in the single-cell data. We then calculate the Pearson Correlation Coefficient (PCC) similarity to find the most similar cell and replace the original spatial cell with this matched single cell. (**F**) Schematic diagram of cell boundary segmentation errors. (**D**) Reconstruction using single-cell and spatial data with simulated segmentation errors. The figure shows the mean Euclidean distance between cells after replacement and their true spatial coordinates, with different colors representing different methods. (**E**) Random sampling of real seqFISH+ data is used to create datasets with cell composition proportions that differ from the original data, to test the impact of noise in cell composition proportions. (**F**) The x-axis shows the cell composition proportions of both the original and sampled datasets, as well as the reconstructed proportions from different methods. The y-axis represents the proportion of each cell type, with different colors indicating different cell types. (**G**) Reconstruction using original seqFISH+ spatial data as single-cell data and sampled spatial data. The figure shows the mean Euclidean distance between cells after replacement and their true spatial coordinates, with different colors representing different methods. (**H**) Different cell types distribution of Ground truth and three reconstructed results.

Our method demonstrated superior performance across multiple evaluation metrics (Supplementary Figure S3B, Figure 2B-2C), achieving the highest spatial distribution similarity and the highest count of true neighbor hits. In terms of spatial distribution similarity of cell types, our method, along with Tangram and CytoSpace, outperformed other methods. Notably, for each specific cell type, our method significantly outperformed Tangram and CytoSpace (Figure 2D, Supplementary Figure S3A). When k=50, the number of true neighbors identified by our method far exceeded that of the other five methods, showcasing Zmap’s ability to accurately reconstruct spatial niche (Figure 2C).

We repeated this test on another spatial profiling technology (STARMAP) using two sections of mouse brain tissue and applied the same methods for mapping. Consistent results were obtained, further validating Zmap’s robustness and precision in spatial reconstruction (Supplementary Figure S4).

### Zmap is robust against spatial data noise

To validate the noise tolerance of Zmap, we simulated a ground truth ST data by substituted each spatial cell of seqFISH+ data by the single cell from snRNA-seq data of the same tissue(17) (Figure 2E). Stringent comparison was performed between Zmap and the state-of-the-art (SOTA) algorithms by integrating the simulated spatial cells and the corresponding single cells. Since the spatial position of each cell was known, we evaluated integration accuracy by computing the average Euclidean distance between mapped and true spatial coordinates of cells. Additionally, we assessed the accuracy of microenvironment reconstruction after mapping by measuring the similarity of spatial distributions across different cell types and evaluating the hit number (true neighbors identified) among the predicted k nearest neighbors.

We tested the impact of cell segmentation noise on integration. In tissues where cells are interconnected, cell boundaries are difficult to distinguish with limited staining, making segmentation errors a common noise source(10). For instance, different algorithms(26,27) recognized macrophage boundaries differently (Supplementary Figure S5A). In some scenarios, even human recognition is difficult to accurately distinguish cell boundaries. To simulate cell segmentation noise, we distributed gene expression from each spatial cell into several parts and randomly allocating them to 2 adjacent cells (Figure 2F). Before perturbation, the UMAP visualization showed clear separation between distinct cell types. However, after perturbation, the intercellular distances decreased, causing different cell types to cluster together (Supplementary Figure S5B). Additionally, each cell type exhibited varying levels of false-positive gene expression (Supplementary Figure S5C). The simulation results were consistent with the recent reports of spatial transcriptomic studies(10). Notably, segmentation noise significantly impaired the performance of Cytospace, and both CelEry and scSpace showed substantial decreases in the number of true neighbors identified. In contrast, Zmap and Tangram were largely unaffected (Figure 2G-2I and Supplementary Figure S5D). We also investigate the impact of imbalanced cell type proportion on the integration (Supplementary Figure S6). These findings demonstrate the advantage of Zmap in resisting cell segmentation noise and highlight its robust adaptability for real-world data integration.

### The versatile application of Zmap on real world data

We now test the accuracy of Zmap by integrating real world datasets. We collected STARMAP spatial sequencing dataset (18) (Figure 3A and 3B) and a snRNA-seq dataset (Figure 3C) (17) from the same cerebral cortex. The two datasets share the same cell types, including excitatory neuron of layer 2-6, inhibitory neurons, astrocytes, oligodendrocytes, etc.

**Figure 3:**
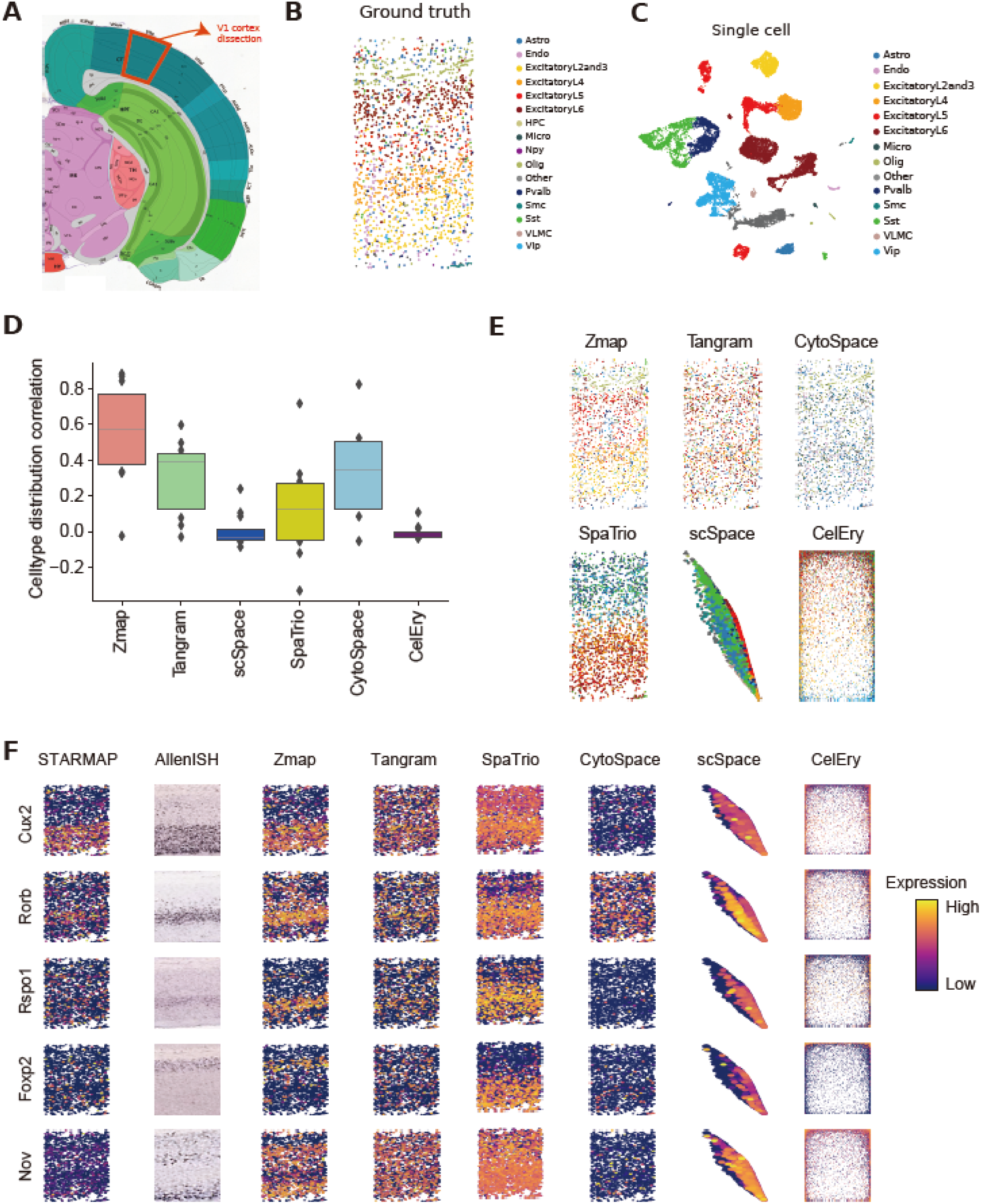
Evaluation in Cerebral Cortex Data. (**A**) Tissue sections from cerebral cortex were used to generate spatial data (STARMAP) and snRNA data. (**B**) Spatial location of main cell types in the STARMAP data. (**C**) UMAP representation of main cell types of the snRNA data. (**D**) Compared cell types spatially reconstructed similarity of six methods. (**E**) Spatial distribution of cell types reconstructed by Zmap and compared methods. (**F**) Reconstructed gene expression.

To evaluate the accuracy of data integration, we measured the spatial distribution patterns of cell types between real spatial transcriptomics data and spatially reconstructed single-cell data. Notably, Zmap shows the highest similarity than other methods, indicating the highest mapping accuracy (Figure 3D). The spatial distribution and proportions of the cell types reconstructed using Zmap resemble the ground truth, as evidenced by the clearly defined hierarchical structures of the cortex (Figure 3E, Supplementary Figure S7 and Supplementary Figure S8B). We also tested the impact of grid size on the results and found that Zmap performed best and outperformed other methods when the average number of cells per bin was between 5 and 14.

Gene coverage is usually limited in ST data due to the probe number in hybridization method or RNA capture efficiency. To test if Zmap can improve spatial gene expression, we used AllenISH tissue staining data as a standard reference (http://atlas.brain-map.org/atlas?atlas=1) (Figure 3F and Supplementary Figure S8A). We first examined the genes common to STARMAP and AllenISH, Zmap successfully predicted the expression of Cux2 in the bottom layer neurons, Rorb in the intermediate layer neurons, and Foxp2 in the top layer neurons.

We also showed that Zmap is powerful in correcting the spatial gene expression. For example, STARMAP failed to present the expression patterns of Rspo1, Foxp2, Nov, and Fezf2, although their probes were designed (Supplementary Figure S8A). Zmap corrected these gene expression and show consistent pattern with AllenISH data. Moreover, Zmap shows ability to recover the gene expression not detected by STARMAP, such as Pvrl3, Agmat, Col5a1 and Cldn11. By such strategy, Zmap increases the spatial genes from 981 (from STARmap data) to 34,041 (from snRNA-seq data). For comparison, the expression pattern reconstructed by other methods was not apparent for all these example genes. These results suggest that accurate cellular localization contributes to more accurate reconstruction of spatial gene expression.

### Zmap enhances the reliability of cell-cell interaction inference

Precision spatial alignment is the prerequisite of accurately identifying intercellular communication, a process critical for delineating proximity-dependent signaling events, understanding the developmental and immune microenvironment for tissue organization. We utilized a seqFISH dataset(14) alongside a scRNA-seq(28) data obtained from the 10x Chromium platform to investigate intercellular communication events in mouse E8.5 embryo (Figure 4A and 4B). The seqFISH dataset detected only 351 genes across 19,416 cells, while the scRNA-seq dataset captured 116,312 cells and 29,452 genes. This significant imbalance between the two datasets introduces noise, including disproportional cell type distribution, posing challenges for spatial integration and complicating the interpretation of intercellular communication. Consistent with the previous analyses, Tangram is more sensitive to uneven cell type distributions compared to Zmap, as the distance between Tangram-mapped single cell coordinates to their nearest spatial cells of the same cell type is much larger (Figure 4C). As displayed by cell type distribution and their correspondence to ground truth, Zmap accurately recapitulates true cell type distribution, whereas Tangram incorrectly predicts a significant portion of cells as erythroid (Figure 4D-4F). The prediction bias generated largely impacts the subsequent ligand-receptor interaction analysis. Erythroid, the cell type with least ligand and receptor counts in the ground truth and Zmap, are considered as the signaling hub by Tangram.

**Figure 4:**
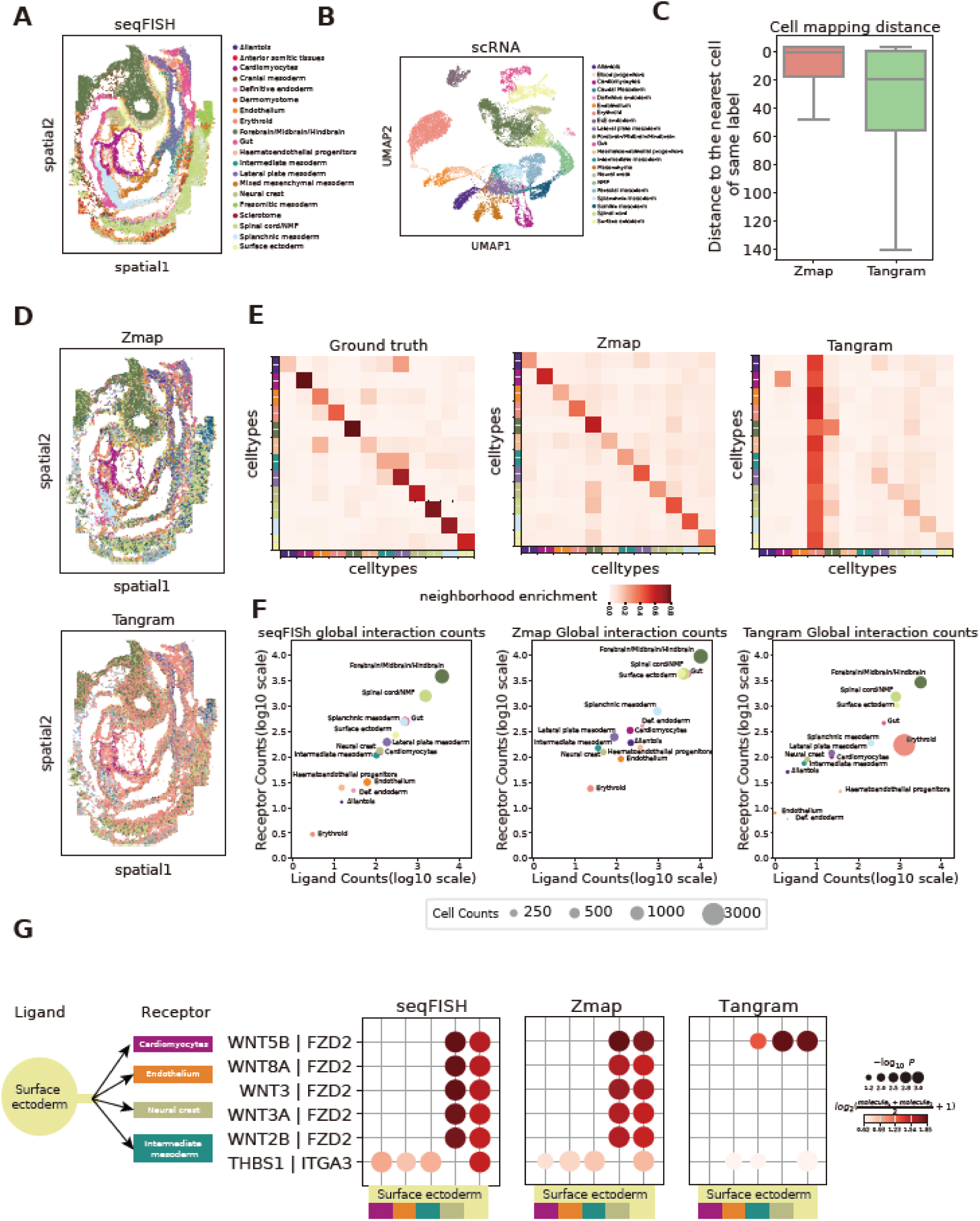
Analysis of Mouse Embryo Data at E8.5. (**A**) Major tissue zones of the mouse embryo at E8.5 using seqFISH. (**B**) UMAP representation of main cell types annotated by Pijuan-Sala et al. (**C**) Compared cell reconstructed distance of Zmap and Tangram; (x axis). We calculated cell reconstructed distances between mapped cells and spatial voxels of the same cell type. (**D**) Spatial distribution of single-cell types after reconstruction by Zmap and Tangram. (**E**) Analysis of spatial neighboring cell proportions of original seqFISH, Zmap and Tangram. (**F**) Number of LR (Ligand-Receptor) pairs analysis with original seqFISH data, Zmap and Tangram reconstructed data, with circle size indicating cell counts. (**G**) LR interactions between surface ectoderm and other cell types.

At E8.5, surface ectoderm serves as a WNT ligand provider for several biological processes, such as neural crest development(29), and optic cup morphogenesis(30). These intercellular communication events are accurately captured by seqFISH and Zmap, showcasing Zmap’s capability to restore precise intracellular communication (Fig. 4g). To exemplify, WNT3A secreted by surface ectoderm induces neural crest fate determination in the presence of FZD2. In contrast, Tangram struggles to identify these ligand-receptor interactions, likely due to spatial mapping errors caused by disproportionate cell types during mismatched data integration.

### Zmap reveals cellular spatial heterogeneity in colorectal cancer-derived liver metastatic tumors

Precise spatial reconstruction is also crucial for tumor microenvironment. Here we applied Zmap to repurpose a 10x Visium data(19) and its corresponding scRNA-seq data of colorectal cancer-derived liver metastatic tumor(19). The spatial data is composed of tumor (T) and para-tumor (PT) regions based on the pathological evidence of the tissue (Figure 5A). Based on the arrangement of 10x Visium spots, we established a regional restriction consisting of four layers: three directional layers at 0 degrees, 60 degrees, and 120 degrees, plus an additional layer that captures the spatial clustering information of the Visium spots for accurate mapping. To assess Zmap’s effectiveness in distributing single cells to these spots, we utilized the Milo algorithm to compute spatial bias scores (SBS) for single cells under T and PT regions. High, low, and medium SBS scores differentiate three single cell statuses—T, PT, and neutrality, respectively (Figure 5B and Supplementary Figure S9A). For instance, the heterogeneous CD8 subpopulations have distinct gene expression profiles (Supplementary Figure S9B), with the upregulated genes in CD8-PT enriching T cell activation terms (Supplementary Figure S9C), indicative of immune activation in PT region.

**Figure 5:**
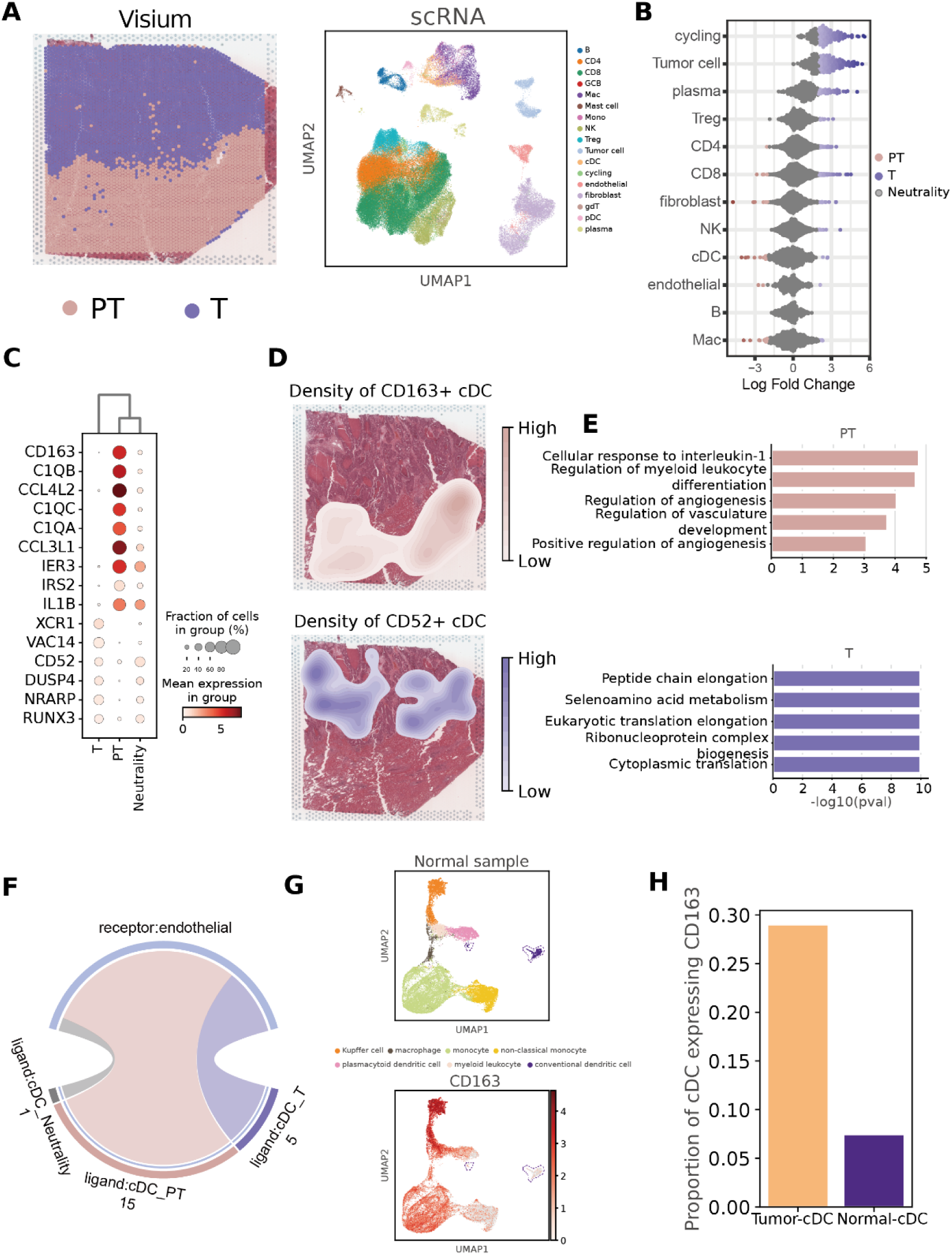
Analysis in Colorectal Cancer Liver Metastasis data. (**A**) Spatial transcriptomics sequencing data (Visium) of colorectal cancer liver metastasis (zones defined by pathology) and UMAP representation of main cell types of scRNA data. (**B**) Spatial distribution preference of different cell types at PT and T zones calculated using Milo. (**C**) Differential genes of cDC (conventional dendritic cell) cell subtypes distributed at different zones. (**D**) Spatial distribution density of different cDC cell subtypes at PT and T zones. (**E**) GO pathway enrichment of differential genes in different cDC cell subtypes, showing that pathways in CD163+ cDC are potentially related to angiogenesis. (**F**) Number of ligand-receptor pairs between endothelial cells and different cDC cell subtypes. (**G**) UMAP representation of healthy liver tissue and the expression of CD163. (**H**) Proportion of CD163 gene expression in cDC cells within liver cancer samples and healthy liver tissue.

Meanwhile, conventional dendritic cells (cDCs) and macrophage contain the most spatial-specific and upregulated genes in PT status (Figure 5B and Supplementary Figure S9A). In recent years, cDC has been found to play an important role in the regulation of immune response and anti-tumor immunity(31,32). However, the spatial heterogeneity and function of cDCs in metastatic colorectal cancer tissues are not clear. CD163+ and CD52+ are defined as differentially expressed genes in the T and PT cDc populations, correspondingly (Figure 5C). The CD163+ and CD52+ density plots reveal the heterogeneity of cDC spatial subtypes (Figure 5D). To further dissect the role of cDC spatial subtypes in regulating immune response, we performed GO enrichment analysis. Genes highly expressed in CD163+ cDC are significantly enriched in angiogenesis pathways (Figure 5E), suggesting CD163+ cDC could promote blood vessel growth and provide nutrients to tumors. The spatial assignment of single cells across cell types (Supplementary Figure S9D) facilitates the identification of cDC spatial subtypes in regulating tumor microenvironment. The ligand-receptor interaction analysis shows that CD163+ cDCs have the highest number of interactions with endothelial cells (Figure 5F). cDC specifically expresses ligand signals related to endothelial genesis, such as NAMPT, SPRED1, PTEN, VEGFA, interacting with endothelial-specific receptors (Supplementary Figure S9E and S9F). Notably, this CD163+ spatial subtype consisting of nearly 30% of cDCs in tumor tissue occupies only 7% of cDCs in healthy liver tissue(33), indicative of its specific role in shaping tumor immune microenvironment (Figure 5G and 5H).

## DISCUSSION

Zmap has a high tolerance for spatial noise, regardless of whether the spatial data is divided into spot, bin or spatial cell. The success of Zmap is attributed to the strategy of cell localization based on multiple regionalization constraints. A region could contain several to hundreds of cells. We claim that the signal-to-noise ratio of a region is higher than that of voxel. This improvement arises because cellular segmentation noise primarily affects the cells in the edge of the region, while the noise among cells within the region tends to cancel out. The joint use of multiple regionalization strategies further enhances the mapping accuracy and the robustness. Consequently, Zmap performed well on datasets from the cerebral cortex, mouse embryos, and tumors.

Zmap offers a flexible framework that allows users to input customized spatial regionalization information, such as spatial clustering outcomes, histological and pathological structures. An increase in the number of layers of spatial regionalization will increase computation time. It is important to note that the division of spatial areas should be meaningful since randomized regionalization may degrade the algorithm’s performance. We suggest using state-of-the-art algorithms to obtain spatial regionalization information as input for Zmap.

As sequencing technology matures and costs decrease, the throughput and resolution of single-cell and spatial data are increasing. Large-scale single-cell atlas studies have become a trend in biological research. When integrating single cell and spatial data, we not only require high accuracy, but also high efficiency of computation. However, existing mapping algorithms cannot efficiently handle millions of cells. Zmap first maps single cells to spatial bins instead of voxels, and the number of cell-to-voxel relationships for calculation is much less than other methods. Therefore, Zmap greatly reduces the cost of computation. At the same time, Zmap uses the pytorch framework for algorithm optimization and supports GPU acceleration to further improve computation efficiency. Zmap provides the ability to reconstruct large-scale integrated atlas for single-cell and spatial data, which would help us to study development, disease, and aging processes from a holistic perspective.

## DATA AVAILABILITY

seqFISH+: spatial transcriptomics data were obtained from https://github.com/CaiGroup/seqFISH-PLUS.

STARMAP data of mouse cortex: scRNA-seq data were obtained from GSE115746, and spatial transcriptomics data were obtained from https://figshare.com/articles/dataset/wang2018three_STATmap/19786456.

STARMAP data of mouse model of Alzheimer’s disease: spatial transcriptomics replicated data were obtained from https://singlecell.broadinstitute.org/single_cell/study/SCP1375

seqFISH: scRNA-seq were obtained from https://github.com/MarioniLab/EmbryoTimecourse2018, and spatial transcriptomics data were obtained from https://marionilab.cruk.cam.ac.uk/SpatialMouseAtlas/.

Visium: scRNA-seq data and spatial transcriptomics data were obtained from GSE225857.

Stereo-seq: spatial transcriptomics data were obtained from https://db.cngb.org/stomics/datasets/STDS0000058

Zmap is also available at https://github.com/JiekaiLab/Zmap

